# *MegaFeed*: Global database of megaherbivores’ feeding preferences

**DOI:** 10.1101/2022.09.23.509174

**Authors:** Fabio Berzaghi, Balram Awasthi

## Abstract

Terrestrial mammalian herbivores heavier than ~1000 kg, also known as megaherbivores, perform unique ecological functions due to their combination of heavy body mass, extended home ranges, abundant biomass consumption, and highly diverse diet. Megaherbivores can have substantial effects on ecosystem functioning, vegetation structure, and biogeochemical cycles. Elephants (family *Elephantidae*) and rhinoceros (family *Rhinocerotidae*) are two of the remaining megaherbivores that survived the late Pleistocene extinctions, but their populations have been globally declining in the last century. Feeding preferences are a key factor determining the influence of megaherbivores on ecosystems and plant communities; however, comprehensive and centralized data on megaherbivores food preferences are lacking. Here we present MegaFeed, an extensive dataset of megaherbivores’ feeding preferences across their distribution. This first version of MegaFeed here described contains more than 12,000 records of feeding preferences for the extant elephant species: *Loxodonta africana* (African savanna elephant)*, Loxodonta cyclotis* (African Forest elephant)*, and Elephas maximus* (Asian elephant). *MegaFeed* will contribute to a better understanding of the ecological functions of megaherbivores, evaluate the consequences of their decline, and guide rewilding and conservation initiatives such as habitat restoration and reduction of human–wildlife conflicts.

## Introduction

Megaherbivores, terrestrial mammals heavier than ~1000 kg, perform important functions in savanna and forest ecosystems and can affect plant species composition, vegetation structure, and biogeochemical cycling (Coverdale et al., 2016; Owen-Smith, 1988; Sitters et al., 2020). Before the Holocene, a wide variety of megaherbivores populated all continents (except Antarctica) and ecosystems. Their extinctions likely triggered important changes in plant diversity, fire regimes, and vegetation cover (Bakker et al., 2016; Gill, 2014; Karp et al., 2021). In the last few millennia, but particularly since the beginning of the industrial revolution, loss of habitat, poaching, and other human disturbances have led to shrinking of extant megaherbivore populations (Ripple et al., 2016). This decline has reduced the scale of megaherbivores ecological functions with repercussions across whole ecosystems (Galetti and Dirzo, 2013; Ripple et al., 2015). Elephants (family *Elephantidae*) and rhinoceros (family *Rhinocerotidae*) are two of the remaining and most studied megaherbivores in Africa and Asia (Berzaghi et al., 2018; Ripple et al., 2015). The critical role that these species play in ecosystem functioning and their global rapid decline make them a priority for research and conservation and for understanding how megaherbivores shaped past ecosystems (Ripple et al., 2016, 2015).

A key ecological trait of megaherbivores is their high daily forage intake coupled with, in some cases, exceptionally broad diets (Owen-Smith, 1988). Elephants and rhinos consume all plant organs (bark, leaf, root, seed, fruit, etc.) of hundreds of different plant species including grasses, trees, shrubs, herbs, and lianas (Blake, 2003; Campos-Arceiz and Blake, 2011; Pradhan et al., 2008). Plant biomass consumption and feeding preferences are thus a main driving force through which megaherbivores shape a variety of ecosystems from grasslands to tropical rainforests (Campos-Arceiz and Blake, 2011; Cardoso et al., 2020; Coverdale et al., 2016; Kortlandt, 1984; Pradhan et al., 2008). A better understanding of the effects of biomass consumption on ecosystems requires in depth knowledge gained from extensive datasets on food preferences. Here we present *MegaFeed*, a global dataset of food preferences of all extant species of elephants (genera *Elephas* and *Loxodonta*) and rhinos (general *Ceratotherium, Diceros, Rhinoceros*, and *Dicerorhinus*). This first version contains data only for elephants, whereas the data collection for rhinos is in its final phase and will be published in the following version. Data in *MegaFeed* cover wide geographical and taxonomical ranges across several countries in Africa and Asia (Fig. 1 and 2). Feeding preferences in *MegaFeed* are reported by plant species, plant functional group, and plant organ consumed (Fig. 2, Tables 1 and 2). When available, data include relative or absolute feeding preference and relative plant abundance at the study site (see Methodology). Additional information includes survey methodology, year of data collection, season, and habitat. *MegaFeed* aims to provide data for biogeography, macroecology, modelling, and phylogenetic approaches aiming at examining the interactions between megaherbivores, plants, and ecosystems with the goal of gaining insights on how these mechanisms influence past and present ecosystem states and composition. *MegaFeed* data can also assist megaherbivore conservation projects involving habitat restoration, reintroductions, and prioritization of conservation areas. Feeding preference data might also help develop strategies to dampen human-megaherbivore conflicts through promotion of less palatable plant species in buffer areas. Large datasets containing information on how food preferences vary through space and time and by species and plant type will be important in developing robust ecological theories and ultimately contribute to more informed conservation decisions.

**Fig. 1.**
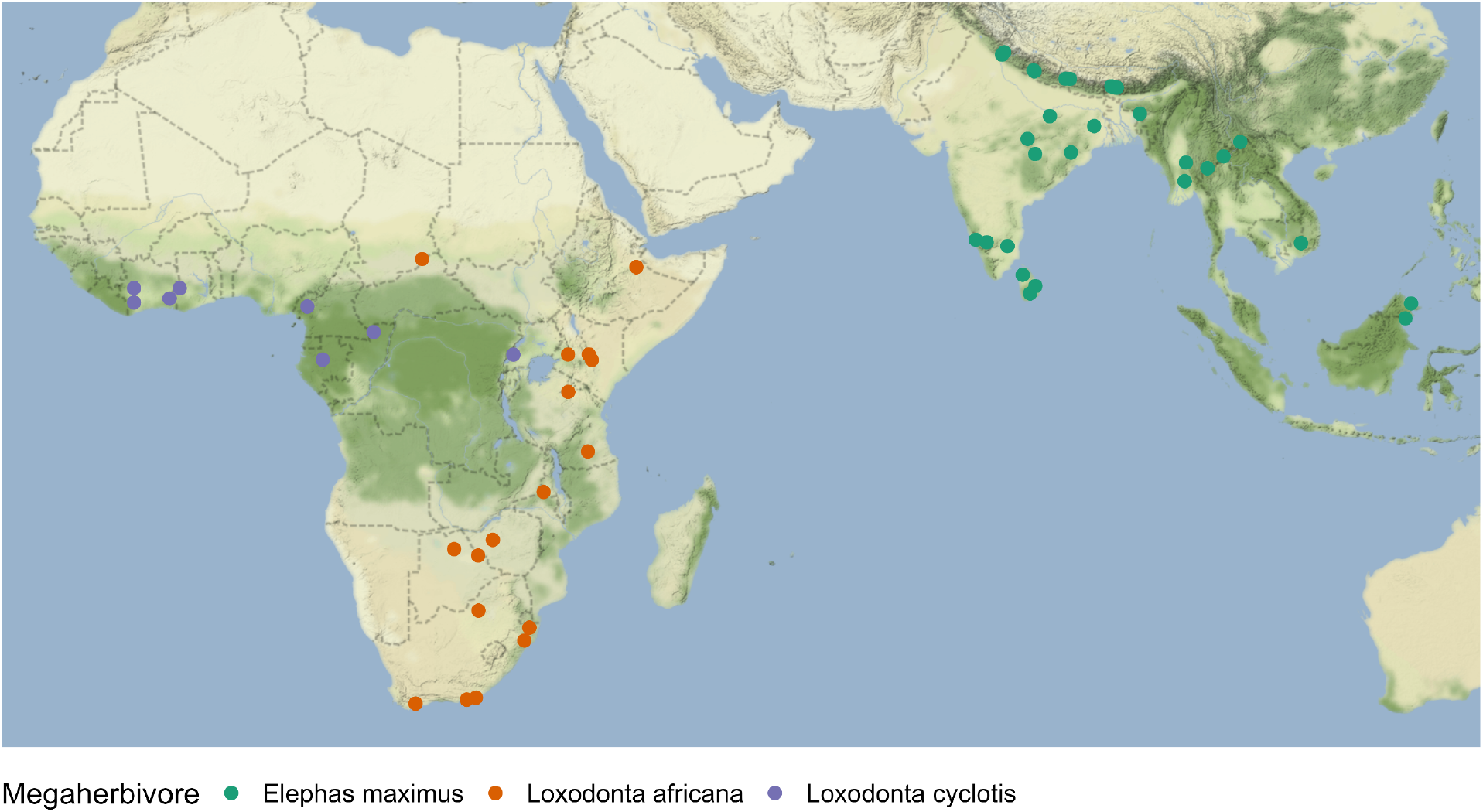
Spatial distribution of data in *MegaFeed* by megaherbivore species. See Table 1 for number of records for each species. In this first version of the database we present data only for the Elephantidae family.

**Fig. 2.**
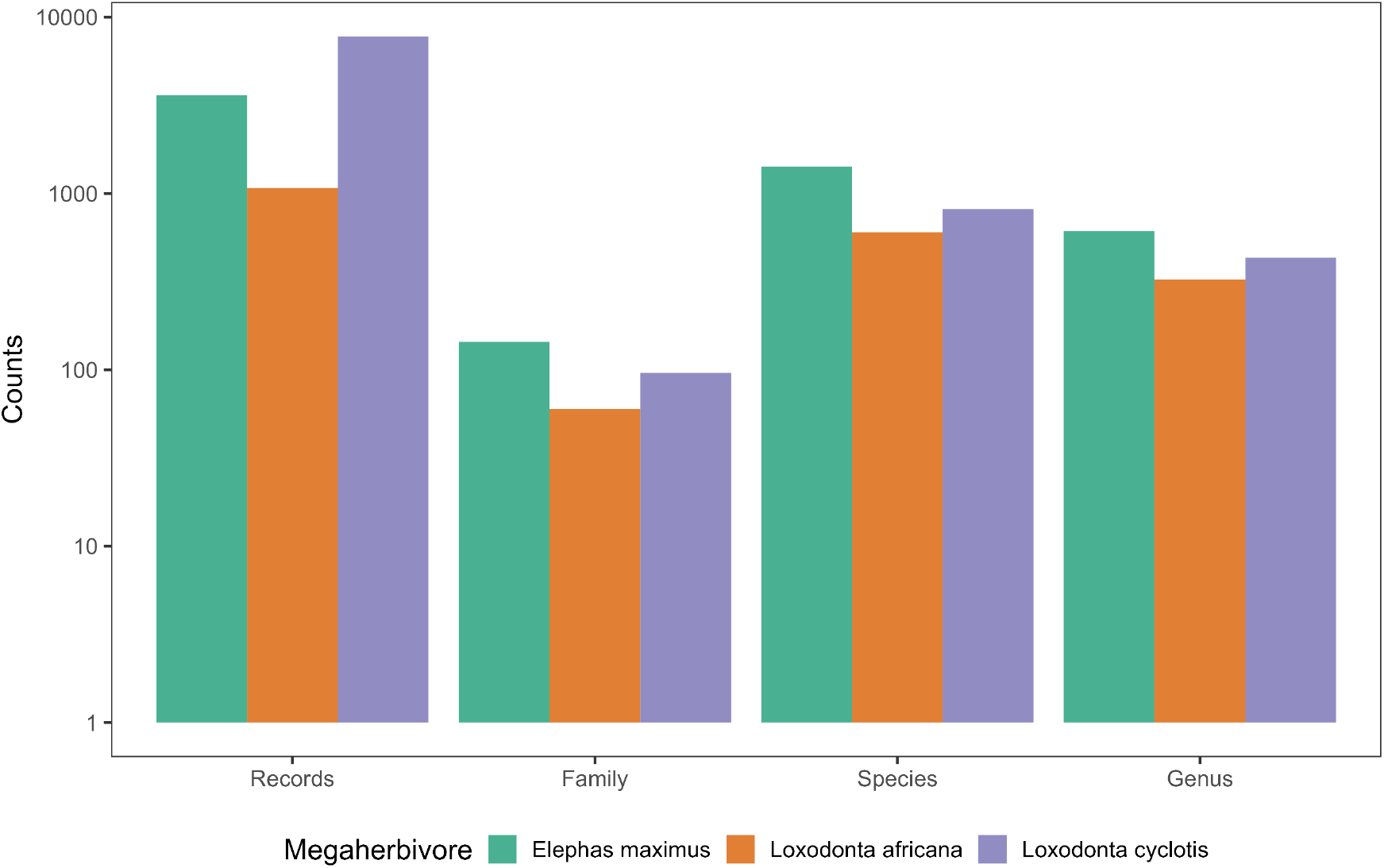
Number of records in *MegaFeed* of (from left to right): feeding records, unique plant families, genera, and species consumed.

**Table 1.** Number of records for which the plant organ consumed by megaherbivores was reported.

| Plant organ eaten | Record count |
| --- | --- |
| leaf | 4197 |
| bark | 1039 |
| leaf+stem | 762 |
| leaf+wood | 437 |
| root | 392 |
| fruit | 389 |
| stem | 219 |
| whole plant | 149 |
| wood | 113 |
| leaf+pith | 85 |
| twig | 81 |
| base | 81 |
| pith | 57 |
| branch | 43 |
| seed | 37 |
| other | 169 |

## Methods

### Data collection

We collected diet of megaherbivores from the literature (peer-reviewed articles and PhD thesis) for Asian elephant (*Elephas maximus*), African savannah elephant (*Loxodonta africana*), African forest elephant (*Loxodonta cyclotis*), white rhino (*Ceratotherium simum simum*), black rhino (*Diceros bicornis*), greater one horned rhino (*Rhinoceros unicornis*), Sumatra rhino (*Dicerorhinus sumatrensis*), and Javan rhino (*Rhinoceros sondaicus*). We performed a literature search using Web of Science and Google scholar with these keywords in a database-style query: “(elephant OR riho* AND feeding preference) OR browsing OR food* OR elephant* OR rhino*”. The same keywords were used in French. We also searched the literature based on our knowledge and the references found in relevant articles. From each paper, we recorded the type of plant and the part consumed, study site (locality, latitude, longitude, country and continent), the year of data collection, the season (when applicable), habitat, and sampling method. When coordinates were not reported, we used OpenStreetMap to estimate latitude and longitude based on the study site description in the article. In most cases, authors reported coordinates for a single point but data collection took place across an area that varied in size. All coordinates were transformed to latitude–longitude (WGS84 decimal degrees). Plant species were updated following the taxonomy of World Flora Online, a global database of plant species. (http://www.worldfloraonline.org/). World Flore Online was also used to determine plant functional type if this was not reported in the article.

In total we found 18 articles for African savannah elephant, 28 articles for Asian elephant, and 10 articles for African forest elephant. We collected a total of more than 12,000 records of feeding preferences that spanned a wide range of plant families, genera, and plant types (Fig. 2 and Table 2).

**Table 2.** Number of records for which the plant type consumed by megaherbivores was reported.

| Plant type | Counts |
| --- | --- |
| tree | 3903 |
| monocot | 1217 |
| liana | 1002 |
| shrub | 588 |
| herb | 461 |
| grass | 363 |
| climber/creeper | 242 |
| browse | 55 |
| bamboo | 47 |
| succulent | 40 |
| crop | 27 |
| fern | 26 |
| forb | 22 |
| tree/shrub | 15 |
| sedge | 6 |

### Description of Data in *MegaFeed*

Most fields in *MegaFeed* are self-explanatory, these include: Megaherbivore, Species (the up-to-date plant species name according to the current taxonomy), Species.original (the original species name reported in the article), Family (the up-to-date plant family according to the current taxonomy), Vernacular.name (plant species name according to the local language), Plant.type (tree, shrub, liana, grass, herb, etc.), Successional.stage (indicates if the plant is a late successional or early/pioneer species and/or shade-tolerant or intolerant; we recorded these definitions only if they were reported in the text), Part.eaten (leaf, seed, fruit, root, pith, bark, branch, twig, flower, stem, etc.), Season (reported only if indicated in the text), Locality, Country, latitude and longitude, Elevation (meters), Habitat, Methods (description of the methodology used to determine feeding preferences), and Author.

#### Feeding preference metrics

The naming of feeding metrics can have some slight variations according to different authors. We grouped feeding metrics in the following categories:

- Percentage.in.diet (indicates the estimated percentage of a particular plant species in the diet of a megaherbivore at the study location)
- Relative.preference (indicates the relative preference of a megaherbivore for a particular plant species at the study location; this can be indicated using a percentage, a ratio, or ordinal categorical units such as low, medium, and high; preference ratios or indices are often calculated by dividing the percentage indicted by the relative abundance, see for examples (Biru and Bekele, 2012; Cardoso et al., 2020)
- Relative.abundance (indicates the relative abundance of a particular plant species at the study location in relation to the abundance of all sampled plant species)
- Number of feeding events (total count of feeding events recorded for a particular plant species at the study location)

## Funding

This work was supported by European Union’s Horizon 2020 research and innovation program under the Marie Sklodowska-Curie grant #845265 and by the French government allocation d’aide au retour à l’emploi program (FB).

## Acknowledgments

We thank all the authors and fieldworkers that collected data in the field and made them available in their manuscripts. Within Stephen Blake for providing feedback on the manuscript.

## Usage notes and data availability

*MegaFeed* data are currently being used for research undergoing peer review. The data will be released gradually as the corresponding research will be published. The dataset will become freely accessible in a spreadsheet format, which will also contain metadata and a complete list of references. An R package will also become available.

## Notes

### Competing Interest Statement

The authors have declared no competing interest.

